# Computer mouse weight is associated with one-week motor skill improvement in a large naturalistic video game cohort

**DOI:** 10.64898/2026.10.06.757150

**Authors:** Jennifer B. Listman, Wayne E. Mackey

**Author notes:** Corresponding author: Jennifer B. Listman.

## Abstract

Computer mouse weight has been proposed to influence human motor performance, but existing evidence comes largely from small, short-duration controlled studies, leaving its relationship to performance during naturalistic gameplay unresolved. We analyzed one-week skill improvement in Aimlabs, a popular first-person shooter aim-training and assessment platform, among 9,067 players whose telemetry recorded a mouse-model change (switchers) during self-directed game use, using a 21-day baseline and 7-day follow-up; 4,991 players with a single mouse model (non-switchers) observed over matched pseudo-anchor intervals provided a secondary contextual comparison. Post-switch mouse weight was represented by eight bands spanning 35-124 g. The primary multivariable generalized additive regression model accounted for baseline skill, pre- and post-switch practice intensity, and pre- and post-switch mouse sensitivity, and allowed the association between weight band and improvement to vary with baseline skill, post-switch practice, and post-switch sensitivity. For each weight band, predicted mean one-week improvement was standardized to the observed distribution of these covariates among switchers and reported in percentile-rank points. Statistical significance was evaluated using 1,000 permutations in which observed mouse weights were randomly reassigned among switchers without replacement and the model was refitted to generate empirical null distributions for the weight-related test statistics. Predicted improvement differed significantly across the eight weight bands (p = 0.001). Pairwise comparisons identified significant differences between bands (p < 0.05), but no single band was associated with significantly greater predicted improvement than all seven others. In this large naturalistic cohort, predicted improvement was highest for players with 35-44 g mice, followed by 45-54 and 55-64 g, and several pairwise comparisons showed significantly greater improvement in lighter than in heavier weight bands. Across the examined baseline-skill profiles, the lowest predicted improvements occurred in either the 85-99 or 115-124 g band. Predicted improvement differed across weight bands at the 25th-percentile and median baseline-skill values (p = 0.002 and p = 0.003) but not at the 75th percentile (p = 0.153). However, the weight-band x baseline-skill interaction was not significant (p = 0.181), indicating that the analysis did not establish that weight-related differences varied by skill level.

## 1. Introduction

Coordinating a limb, or a hand-held tool, to produce a precise, goal-directed movement requires resolving many redundant mechanical degrees of freedom. This problem was posed classically by Bernstein (1967) and reframed in modern computational terms as a question of which internal representations of the body and the world the motor system must learn and update (Wolpert and Ghahramani, 2000). Motor learning is typically studied in the laboratory, where the task is simple, the environment is controlled, and the equipment a participant uses is fixed by the experimenter (Fitts and Posner, 1979; Krakauer et al., 2019). Large-scale, naturalistic studies relax these constraints, trading experimental control for sample size, task complexity, and ecological validity.

Computer-based video games provide an unusually rich setting in which to study complex motor behavior over time. A study by Stafford and Dewar (2014) used online game data from over 800,000 participants to confirm previously experimentally validated laws of skill acquisition such as the relationships between practice amount or practice spacing and subsequent performance. Methodological work has argued that large naturalistic samples can complement small laboratory cohorts by improving generalizability (Hilton and Mehr, 2022). Online sensorimotor experiments have also reproduced established adaptation effects using participants’ own computers and input devices (Tsay et al., 2021). The present study uses intrinsically motivated video game play observed at scale to examine whether mouse weight is associated with motor performance.

Aimlabs (Statespace Labs, Inc., New York, NY, USA; https://aimlabs.com/) is a commercial software platform for assessing and training visuomotor skills used in first-person shooter games. In this genre, a player views a three-dimensional virtual environment from the character’s perspective and uses a mouse to rotate the view, aim a virtual weapon, and select or shoot visual targets. Aimlabs provides task scenarios designed to assess and train components of this performance, including rapid target acquisition, aiming precision, and tracking of moving targets. Each task produces a score, additional performance metrics, and associated gameplay telemetry.

Aimlabs first person shooter training tasks have been used in studies of motor neuroscience and to interrogate human cognitive and motor performance change under varied training, physiological, and exercise conditions. Keeler et al. (2023) used Aimlabs tasks to test the effects of moderate hypoxia on cognitive performance and Wachholz et al. (2025) used Aimlabs tasks to test the effects of physical exercise on gaming performance. Aimlabs task data has also been used to study long-term motor learning (Listman et al., 2021), aiming expertise and movement kinematics (Donovan et al., 2022), and attentional focus (Lamers James and O’Connor, 2023).

In previous work by our group, Aimlabs players selected their own tasks, practice schedules, and duration of play, producing a naturalistic record of practice under heterogeneous, uncontrolled conditions (Listman et al., 2021). Across more than 7,000 players, motor acuity improved steadily over as many as 100 days, and learning rate varied with baseline skill, daily practice volume, and the interval between sessions. Despite variation in players’ equipment and environments and many unknown participant-level covariates, the analysis reproduced several well-established findings from controlled motor-learning studies. This convergence supports the use of Aimlabs telemetry as an instrument for large-scale motor-neuroscience research in naturalistic settings (Listman et al., 2021).

Practice-driven learning and adjustment to a changed device can occur at the same time. Aimlabs lets a player set mouse sensitivity, which determines the view rotation produced by a given physical displacement (Donovan et al., 2022; Boudaoud et al., 2023). A different mouse can also change the physical dynamics of aiming through its mass and mass distribution, even when software sensitivity remains unchanged (Baek et al., 2025). Fast recalibration to a changed sensorimotor mapping is usually described as motor adaptation, whereas the refinement of a skill through extended practice occurs over longer time scales (Haith and Krakauer, 2013; Krakauer et al., 2019; Listman et al., 2021). The present observational data cannot isolate these mechanisms, but they can estimate the conditional association between mouse weight and performance improvement while accounting for measured practice volume, player skill, and mouse sensitivity. Tsay et al. (2024) demonstrated how large-scale citizen-science data can be used to identify predictors of sensorimotor adaptation. Their analysis included broad device type (mouse, trackpad, or trackball) among many candidate predictors. The present study complements that work by resolving mass within one device class.

The mouse itself has been studied in human-computer interaction for decades, principally through work on pointing-device geometry and control mapping and additional studies have shown that pointing performance depends jointly on the user, the control mapping, and the physical device. Early comparisons showed that pointing-device choice affects target-acquisition speed and error rate (Card et al., 1978; MacKenzie et al., 1991). More recent online experiments found that mouse and trackpad input produced different reaction times and movement kinematics, further demonstrating that the physical input device can affect measured behavior (Warburton et al., 2025). Fitts (1954) formalized the relationship between movement time, target distance, and target width; Fitts-law throughput subsequently became a standard measure for evaluating pointing devices (Douglas et al., 1999; Soukoreff and MacKenzie, 2004; Zhai et al., 2004). Even when the physical device is held constant, control-display gain can affect pointing performance (Casiez et al., 2008; Sandfeld and Jensen, 2005). In first-person perspective targeting, mouse sensitivity similarly determines the mapping between hand displacement and view rotation (Boudaoud et al., 2023).

Controlled studies have tested effects of mouse mass on target acquisition or video-gaming performance directly. These studies typically enroll tens of participants and generally assessed performance during brief laboratory sessions, with findings that differed by outcome and weight range. Studies of professional or high-level gamers using mice that differed by 7-13 g found preferences for lighter options but little difference in several performance, muscle-activity, or kinematic outcomes (Li et al., 2019, 2022; Wang et al., 2023). A study spanning 40-140 g found the highest error rate with the lightest mouse tested and the highest subjective preference near 60 g (Yan et al., 2022), whereas a point-and-click study spanning 70-190 g found a non-monotonic association between weight and forearm muscular load (Chen et al., 2012). Moment of inertia and center-of-mass distribution may also affect perceived mouse weight independently of gram count (Baek et al., 2025). Conroy et al. (2022) reported faster and more accurate target acquisition with 50-90 g mice than with a 100 g mouse in an action-video-game task. While informative about immediate performance, preference, and muscle activity, these studies do not establish whether mouse weight is associated with performance change over a longer period.

Motor adaptation to changes in implement mass has also been studied outside computer use. Batters recalibrated their movements using strategies specific to lighter vs heavier bat weight (Scott and Gray, 2010), and the response to a change in badminton-racket weight differed between advanced and beginning players (Pan et al., 2024). In a simulated batting task, switching to either a lighter or heavier bat initially increased swing-timing errors, but participants recalibrated within 5-10 trials by adjusting swing velocity for the lighter bat and swing-onset timing for the heavier bat (Scott and Gray, 2010). In badminton jump smashes, advanced players adapted to a heavier racket primarily by changing the timing and coordination of muscle activation, whereas beginners relied more heavily on changes in upper-limb muscle activity, suggesting that motor experience permits faster and more coordinated adaptation to increased implement weight (Pan et al., 2024).

In the present study, the primary analysis examined one-week change in cross-task calibrated percentile rank among switchers after an observed mouse-model change, such that the incoming hardware exposed them to a change in mouse weight. Players who did not change their mouse were retained for a secondary contextual comparison rather than as a conventional control group. Because commercially produced gaming mice are not distributed evenly across the weight range, post-switch weight was represented by eight bands defined from the observed exposure support. A multivariable generalized additive regression model included baseline skill, pre- and post-anchor practice intensity, and pre- and post-anchor mouse sensitivity, with interactions allowing the association between weight band and improvement to vary with baseline skill, post-anchor practice, and post-anchor sensitivity. We asked whether predicted one-week improvement differed across the eight weight bands among switchers and whether the complete weight-band pattern varied with baseline skill. We subsequently compared the switcher results with predictions from non-switchers observed over matched no-change intervals. Because device weights and mouse-model changes were observed rather than assigned or randomized, the results are interpreted as associations rather than causal effects.

## 2. Methods

### 2.1 Study design and observation windows

This was a retrospective observational analysis of de-identified Aimlabs gameplay and device telemetry data collected during the course of typical, self-directed Aimlabs sessions. Mouse use and any mouse-model changes occurred outside investigator control; no mouse manufacturer or model, mouse weight, practice schedule, Aimlabs task selection, or mouse sensitivity setting was assigned by the investigators. Players whose telemetry recorded a change from one identified mouse model to another were classified as switchers. Players whose telemetry showed stable use of one identified mouse model during an eligible comparison interval were classified as non-switchers. For a switcher, the first date on which telemetry showed an incoming mouse model different from the previously recorded model became the anchor date. This was the first observed calendar day on which at least one eligible Aimlabs play showed a mouse model different from that previously detected for a given player; it is possible that the player used this mouse previously outside of Aimlabs so this date is not necessarily the player’s first physical use of it. When a player had more than one observed mouse change with an eligible 21-day baseline and seven-day follow-up window, one change and its associated data were randomly selected so that each player contributed one switcher observation. Non-switchers were assigned a pseudo-anchor date sampled from a period of stable use of a single recorded mouse model. A skill calibration population of 7,001,710 unique users was selected from data collected through Aimlabs play between May 27th, 2025 through May 27th, 2026.

A player’s baseline skill window comprised the 21 days before the anchor date. The post-anchor window comprised days +1 through +7. Player performance data on the anchor day was excluded from analysis because it could represent a partial exposure day containing Aimlabs plays made with both the outgoing and incoming mouse. Changes in equipment or control settings can also produce transient adaptation-related performance disruption (Smith et al., 2006; Krakauer et al., 2019). In unpublished analyses of a large Aimlabs cohort, our research group similarly observed a temporary performance decrease on the day of a mouse-sensitivity change that recovered on the next day of play.

Non-switcher candidates had at least three consecutive calendar months of recorded use of the same mouse and at least five plays per month in the upstream monthly table. Each non-switcher was assigned a no-change pseudo-anchor date within that stable-use period. The date was required to leave the complete 21-day baseline and seven-day follow-up windows within the same-mouse run, with an additional seven-day buffer between the analysis windows and each recorded run boundary. Up to 1,500 non-switchers were sampled per predefined weight bucket, and their pseudo-anchor-week distribution was matched to the switch-week distribution. Thus, the non-switcher cohort supplied a matched-time comparison during stable mouse use; its raw weight distribution reflects the sampling design applied to switchers rather than mouse weight prevalence among Aimlabs players.

Mouse manufacturer and model identity was resolved before construction of the analysis cohort. The device-attribution pipeline combined internationally standardized vendor and product identifiers with device strings collected through Aimlabs standard telemetry to assign a canonical manufacturer and exact model. Switchers were eligible only when both the outgoing and incoming mouse models were identified; non-switchers required an identified model that remained unchanged throughout the relevant stable-use period. Each mouse model also had to have a publicly documented weight in grams. The exposure value was the manufacturer-listed device-only weight; cables, receivers, and other external accessories were not included. Reliable mouse identity was therefore used to establish eligibility, switching status, and weight, but manufacturer and model identity were not included as variables in the statistical model nor to design the sample.

### 2.2 Performance, sensitivity, and participant eligibility

A play constituted one completed, eligible 60-second Aimlabs task that trained and assessed skills used in first-person shooter video games. Duplicate, aborted, and otherwise ineligible events, including extreme score values indicative of error, cheating, or another anomaly, were removed upstream. The primary cohort required at least five total plays across the pooled 21-day baseline period and at least five plays during days +1 through +7; five plays were not required in each baseline week. A sensitivity-linked play was an eligible play with valid recorded horizontal sensitivity (sens_x) and DPI, permitting calculation of horizontal effective DPI (eDPI = sens_x x DPI). At least two sensitivity-linked plays were required in each period. When settings varied within a period, eDPI was calculated for each play and the period value was the arithmetic mean across sensitivity-linked plays; equivalently, weekly summaries were weighted by their numbers of sensitivity-linked plays. Aimlabs’ calibration is 0.05 degrees per mouse count, so degrees per inch = 0.05 x eDPI and cm/360 = 18,288/eDPI (Donovan et al., 2022). The constant 18,288 equals 360 degrees x 2.54 cm per inch divided by 0.05 degrees per count.

An outcome-independent telemetry-validity screen required DPI between 50 and 50,000, derived sensitivity between 1 and 1,000 cm/360, no more than a ten-fold pre/post eDPI change, and at least two sensitivity-linked plays in each period. The same rules were applied to switchers and non-switchers. These bounds were intended to remove implausible telemetry rather than unusual but physically possible player settings.

Some participants appeared once as a switcher and once as a sampled non-switcher in non-overlapping time frames. To retain one independent observation per player, if one cohort-by-weight cell contained fewer than 30 observations and the corresponding cell in the other cohort was at least twice as large, the player was assigned to the smaller cell to preserve sparse exposure support. The remaining duplicated players were assigned to a single cohort by a reproducible 50:50 draw.

The data set was then restricted to post-anchor mouse weights from 35 through 124 g, the supported range in which every weight band (section 2.3) contained at least 30 participants from each cohort yielding 14,058 unique participants: 9,067 switchers in the primary analysis and 4,991 non-switchers in the secondary comparison.

### 2.3 Mouse-weight exposure

The exposure was the post-anchor mouse weight obtained from the exact-model lookup described above. For switchers, this was the incoming mouse weight; for non-switchers, it was the weight of the continuously used mouse. Inference was restricted to 35-124 g, where every weight band contained at least 30 participants from each cohort. Before outcome differences were inspected, the following eight bands were fixed from the observed exposure-support distribution: 35-44, 45-54, 55-64, 65-74, 75-84, 85-99, 100-114, and 115-124 g. The narrower 10-g bands cover the denser lower range; the upper bands are 15 g wide where exposure support was sparser. The boundaries were not produced by an outcome-optimized selection algorithm.

### 2.4 Skill calculation and outcome construction

There are over one thousand Aimlabs tasks that assess and train skills used in first-person shooter video games, and tasks differ in both scoring scale and popularity. To place eligible plays on a common 0-1 skill scale, player percentile rank for a given task was calibrated against their rank on four high-volume anchor tasks, each with 1.3 million - 2.7 million plays from the general Aimlabs player population over the course of the data observation period: Gridshot Ultimate, Sixshot Ultimate, Valorant Small Flicks, and VT Miniphase VALORANT. For each player-task pair in the fixed study-year calibration population, the player’s best score in any of these four tasks was converted to a percentile rank. Non-anchor task scores were divided into 20 score bins. For each task-bin-anchor combination, the mean anchor-task percentile was calculated among players who had performed both tasks; a task-anchor mapping required at least five shared players per bin and at least 50 shared players across bins. Each anchor-specific curve was constrained to be nondecreasing by a cumulative-maximum operation. Qualifying anchor curves were then averaged using their numbers of shared players as weights, and the combined mapping was again constrained to be nondecreasing. Anchor-task plays retained their direct percentile ranks. Non-anchor plays without a qualifying cross-task mapping did not contribute to baseline skill, post-anchor skill, or improvement but counted toward practice intensity. Scores from over seven million unique Aimlabs players contributed to the study population skill calibration calculations.

For participant *i*, baseline skill was the mean calibrated rank across all eligible plays during the three baseline weeks. Post-anchor skill was the mean calibrated rank across eligible plays on days +1 through +7. The outcome, improvement, was calculated as post-anchor skill minus baseline skill. Thus, positive values indicate higher calibrated performance during the post-anchor week. Baseline skill remained referenced to the broader Aimlabs calibration population (N > 7 million) and was not re-ranked within the switcher and non-switcher study cohort. Calibrated rank ranges from 0 to 1; figures and tables multiply it by 100 for readability.

### 2.5 Covariates and statistical analysis

The primary analysis was restricted to the 9,067 switchers in the supported 35-124 g range who had complete pre- and post-anchor skill calibration. A Gaussian generalized additive regression model with an identity link was fitted by restricted maximum likelihood using mgcv in R (Wood, 2011). Weight was represented by the eight-level categorical factor described above. Baseline skill, post-anchor practice intensity, and post-anchor mouse sensitivity were each represented by a natural cubic spline with three degrees of freedom, and each spline was interacted with weight band. Three degrees of freedom were specified before fitting as a parsimonious allowance for curvature: this permits departures from a straight-line association while limiting instability in the smaller weight-band cells.

The model estimated one-week calibrated-rank improvement for each mouse-weight band, potentially nonlinear associations with baseline skill, post-anchor practice intensity, and post-anchor mouse sensitivity, and the interaction of each association with weight band. It also included a smooth interaction between post-anchor practice and baseline skill, plus separate smooth adjustment terms for pre-anchor practice and pre-anchor mouse sensitivity. Practice volume was transformed as the natural logarithm of one plus the number of plays because our earlier Aimlabs motor-learning analysis found a nonlinear association between practice volume and improvement, with diminishing returns as within-day practice increased (Listman et al., 2021). Mouse sensitivity was natural-log transformed because eDPI is directly proportional to view rotation per unit of physical mouse movement; the log scale represents equal proportional changes equally across the sensitivity range and reduces the influence of extreme values.

The weight-band interactions with baseline skill, post-anchor practice intensity, and post-anchor mouse sensitivity were represented by natural cubic spline functions, each with three fixed degrees of freedom. An additional penalized smooth term modeled the joint nonlinear association of post-anchor practice and baseline skill beyond their separate associations. Pre-anchor practice and pre-anchor mouse sensitivity were included as separate penalized smooth terms. Smoothing parameters were estimated by restricted maximum likelihood, with shrinkage allowing weakly supported smooth components to approach zero. In the fitted model, the effective degrees of freedom were 4.20 for the post-anchor practice-by-baseline-skill interaction, 1.14 for pre-anchor practice, and 0.92 for pre-anchor mouse sensitivity.

For each weight band, covariate-standardized predicted improvement was calculated by assigning that same band to every switcher in the model while retaining each player’s observed baseline skill, practice, and sensitivity values, and then averaging the 9,067 player-level predictions. This produces the predicted mean one-week percentile-rank improvement for a common switcher covariate distribution, enabling comparisons among weight bands that are not driven by measured differences in those covariates. All predicted improvements and contrasts are reported as percentile-rank points, obtained by multiplying the 0-1 calibrated-rank outcome by 100. Pointwise 95% confidence intervals were calculated as the prediction plus or minus 1.96 standard errors, using a covariance matrix that included smoothing-parameter uncertainty. An omnibus test evaluated whether the eight band predictions were equal, and a separate interaction test evaluated whether the complete weight-band pattern changed with baseline skill. Predictions were also calculated at the 25th percentile, median, and 75th percentile of baseline skill while standardizing the remaining covariates over the switcher distribution.

Because the covariate-standardized predictions were derived from the same model and were statistically correlated, overlap between their pointwise 95% confidence intervals was not used to determine whether weight bands differed. Differences between bands were evaluated using covariance-aware pairwise contrasts, with statistical significance determined from the corresponding raw empirical permutation p values. The 28 overall comparisons and the 28 comparisons at each of the 25th-percentile, median, and 75th-percentile skill profiles were evaluated; all comparisons with raw empirical p < 0.05 are reported together so that the complete pattern can be assessed (Table 3).

In addition to the switcher-only primary model, a separate joint model of switchers and non-switchers was used as a secondary, post hoc contextual comparison, not as a randomized or conventional control analysis. The joint model used the same eight weight bands and covariate classes and added interactions with cohort. Its three related omnibus tests (differences across weight bands among non-switchers, differences across weight bands among switchers, and differences between the complete switcher and non-switcher weight-band patterns) were treated as one family and evaluated with Holm correction for multiple testing (Holm, 1979). Model review included convergence, residual scale and fit summaries, and saved basis-dimension and concurvity diagnostics. The primary switcher-only model converged with residual scale < 0.005, adjusted R-squared 0.147, and 15.5% deviance explained.

### 2.6 Mouse-weight-label permutation analysis

Because the sampling distributions of the model-based test statistics were not assumed to follow a known parametric form, statistical significance was evaluated using permutation tests. Empirical null distributions were generated by randomly reassigning the observed mouse-weight labels among players without replacement, thereby preserving the number of players in each weight band, and refitting the regression model to each permuted dataset (Nichols and Holmes, 2002; Pitman, 1937; Winkler et al., 2014). In each permutation, the 9,067 observed post-switch mouse weights were randomly reassigned among the 9,067 switchers without replacement. Thus, every observed weight was used exactly once in each permuted data set and the number of players in each weight band was preserved, while the outcome and all non-weight covariates remained attached to their original players. The eight weight bands were reconstructed from the reassigned weights, and the same generalized additive regression model was refitted by restricted maximum likelihood, including re-estimation of smoothing parameters. The weight-related omnibus, interaction, profile-specific, and pairwise statistics were saved after each fit. This process was completed 1,000 times, producing a separate null distribution for every reported weight-related statistic; all 1,000 models converged.

For each test, the statistic from the observed data was compared with its corresponding distribution from the 1,000 permuted data sets. The raw empirical p value was calculated as (1 + the number of permuted statistics at least as extreme as the observed statistic)/1,001. The added 1 in the numerator and denominator prevents a p value of zero when none of the 1,000 permuted statistics is as extreme as the observed value. Omnibus tests evaluated differences in average predicted improvement across the eight weight bands, all weight-related model terms jointly, the interactions of weight band with baseline skill, post-anchor practice, and post-anchor sensitivity, and differences across bands at the 25th percentile, median, and 75th percentile of baseline skill. Pairwise tests were two-sided and compared the absolute observed contrast statistic with its corresponding permutation distribution. With 1,000 permutations, the smallest attainable empirical p value was 0.001.

Because mouse weight was observed rather than experimentally assigned, the weight-label permutation procedure was used as a sensitivity analysis rather than interpreted as random assignment of exposure. It tests whether the observed weight-related statistics are larger than those produced after randomly breaking the observed player-to-weight assignment; it does not remove unmeasured confounding or establish that mouse weight caused the observed differences.

### 2.7 Ethics, data availability, and software

Ethical considerations: This study analyzed strictly anonymized, retrospective telemetry data automatically generated during routine gameplay. The dataset contained no personally identifiable information, demographics, or tracking mechanisms that could link data to individual players. Because the research involved only secondary analysis of fully de-identified commercial data, with no direct interaction with users and no experimental manipulation of gameplay, it does not constitute human-subjects research under federal guidelines (e.g., 45 CFR 46), and formal institutional review board approval was not required. In compliance with the platform’s terms and privacy rules, all users must be aged 13 years or older to play.

The proprietary row-level telemetry cannot be made public because it was collected for commercial operations and would disclose details of Aimlabs data systems. Requests for access should be directed to and are subject to company approval and applicable privacy restrictions. Analysis code, model specifications, diagnostics, and non-identifying derived results used to reproduce the reported tables and figures will be deposited in a public repository; a permanent link will be added to a subsequent version of the manuscript.

## 3. Results

### 3.1 Analysis cohort and exposure support

Cohort construction began with 30,344 candidate participant-window records representing 29,659 unique players across the switcher and non-switcher sampling procedures. Requiring at least five baseline and five post-anchor plays over a single 29-day window retained 17,056 records representing 16,778 players. Complete pre- and post-anchor mouse-sensitivity telemetry retained 15,545 records. Application of the outcome-independent telemetry-validity rules retained 15,528 records representing 15,272 players. After assigning each overlapping player to one cohort and restricting post-anchor weight to the supported 35-124 g range, 15,098 participants remained. Requiring complete cross-task calibration excluded 1,040 participants, leaving 14,058 participants: 9,067 switchers and 4,991 non-switchers. Every band included at least 224 switchers and 285 non-switchers, although support was uneven (Table 1). Switchers were concentrated in the 45-64 g range, whereas the quota-sampled non-switcher distribution was flatter and should not be interpreted as population prevalence.

**Table 1.** Sample size (n) by post-anchor mouse-weight band in the mouse switcher and non-switcher cohorts.

| Mouse weight band (g) | Non-switchers, n | Switchers, n |
| --- | --- | --- |
| 35-44 | 599 | 503 |
| 45-54 | 654 | 3,481 |
| 55-64 | 637 | 3,313 |
| 65-74 | 651 | 303 |
| 75-84 | 615 | 315 |
| 85-99 | 1,074 | 604 |
| 100-114 | 285 | 324 |
| 115-124 | 476 | 224 |

All eight mouse weight bands were represented in both cohorts. Switcher counts reflect observed post-switch mouse weight. Non-switcher counts reflect quota sampling.

### 3.2 Primary mouse switcher-only analysis

The primary switcher-only model included 9,067 switchers and converged with 119 coefficients and 86.26 total effective degrees of freedom. Adjusted R-squared was 0.147, deviance explained was 15.5%, and the residual scale estimate was 0.005449. Covariate-standardized predicted one-week improvement differed significantly across the eight post-switch weight bands (raw empirical permutation p = 0.001). The direct weight-band x baseline-skill interaction was not statistically significant (p = 0.181). Thus, the analysis supports an overall association between weight band and predicted improvement but does not show that the complete association differs by baseline skill.

The largest predicted one-week improvement was 2.74 percentile-rank points (95% CI, 2.05 to 3.42) for 35-44 g, followed by 2.58 points (2.34 to 2.82) for 45-54 g and 2.10 points (1.87 to 2.34) for 55-64 g. Predictions were 2.06 points for 65-74 g, 1.70 for 75-84 g, 0.67 for 85-99 g, 1.80 for 100-114 g, and 0.52 for 115-124 g (Figure 1; Table 2). The higher predicted improvement at 100-114 g than at 85-99 or 115-124 g shows that the categorical pattern was not monotonic.

**Table 2.** Covariate-standardized predicted one-week percentile-rank improvement among switchers.

| Weight band (g) | Switchers, n | Covariate-standardized predicted improvement (percentile-rank points) | Pointwise 95% CI (percentile-rank points) |
| --- | --- | --- | --- |
| 35-44 | 503 | 2.74 | 2.05 to 3.42 |
| 45-54 | 3,481 | 2.58 | 2.34 to 2.82 |
| 55-64 | 3,313 | 2.10 | 1.87 to 2.34 |
| 65-74 | 303 | 2.06 | 1.27 to 2.85 |
| 75-84 | 315 | 1.70 | 0.89 to 2.50 |
| 85-99 | 604 | 0.67 | 0.06 to 1.28 |
| 100-114 | 324 | 1.80 | 0.95 to 2.64 |
| 115-124 | 224 | 0.52 | -0.50 to 1.55 |

*For each weight band, predictions were averaged across the observed covariate values of all 9,067 switchers as described in section 2.5. Confidence intervals are pointwise 95% intervals. All values are percentile-rank points (the 0-1 calibrated-rank outcome multiplied by 100).*

**Figure 1.**
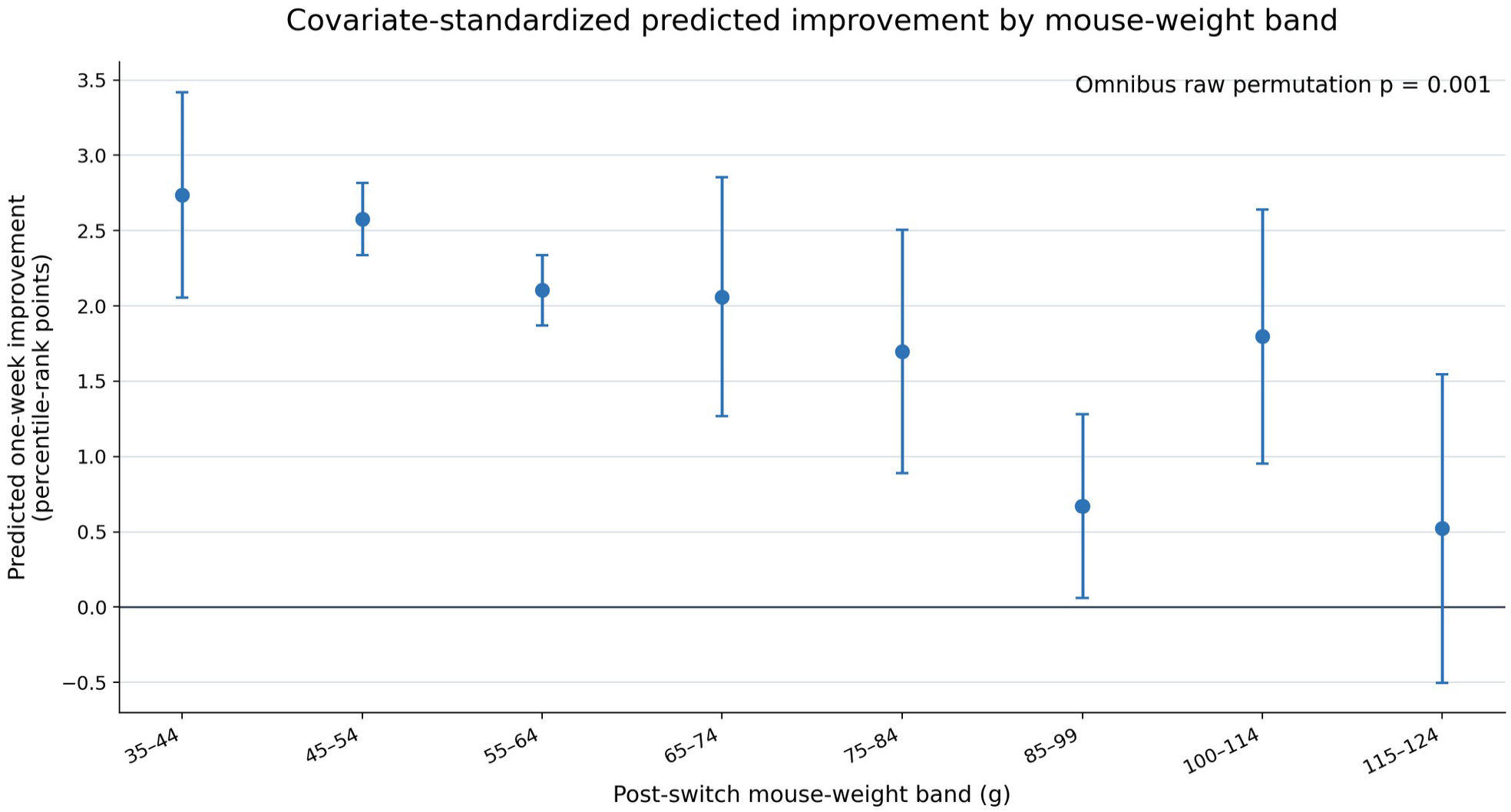
Covariate-standardized predicted one-week percentile-rank improvement by post-switch mouse-weight band among switchers. For each band, points show the average model prediction across all 9,067 switchers after assigning that band to every player while retaining each player’s observed covariates; error bars show pointwise 95% confidence intervals. Values are percentile-rank points. The displayed p value is the raw empirical p value from the 1,000 weight-label permutations for the omnibus test of equality across all eight bands. Because the predictions are statistically correlated, overlap of their confidence intervals is not a test of between-band differences.

**Figure 2.**
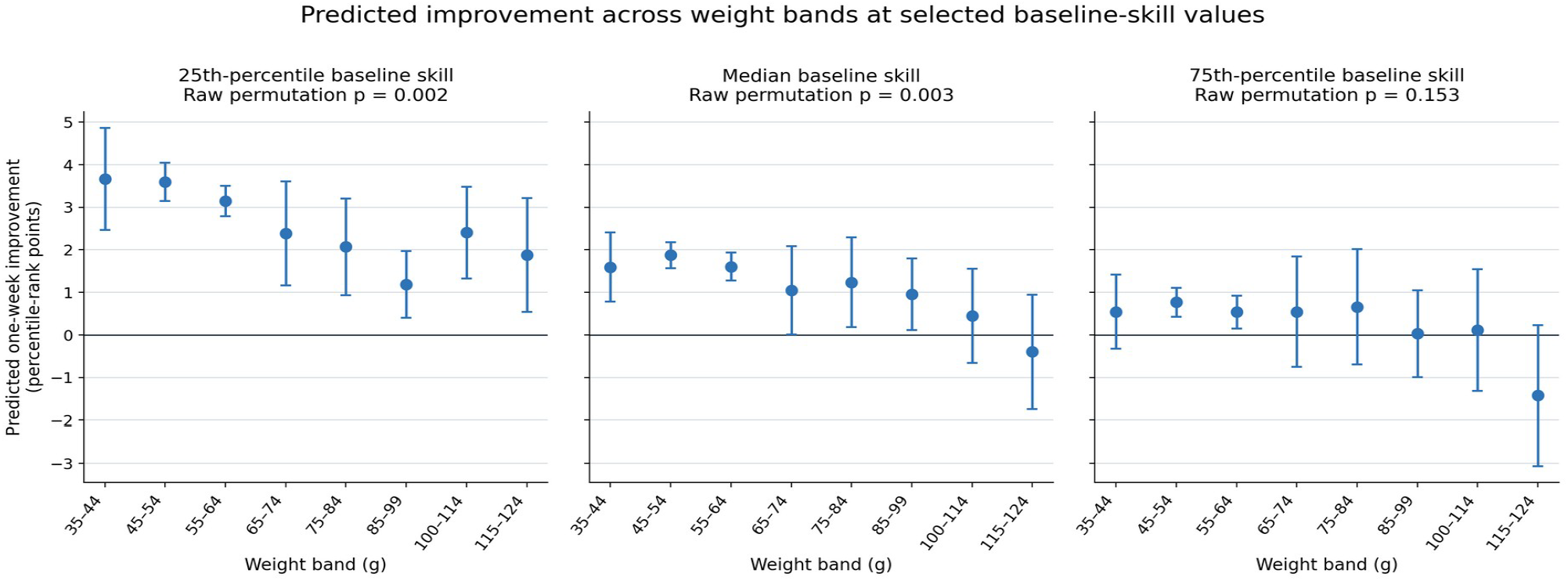
Predicted one-week percentile-rank improvement across all eight post-switch mouse-weight bands at the 25th percentile, median, and 75th percentile of baseline skill among switchers. Baseline skill was fixed at the displayed value while the remaining covariates were standardized over the switcher distribution. Points show predicted improvement and error bars show pointwise 95% confidence intervals. Panel p values are raw empirical values from the 1,000 weight-label permutations for the omnibus test of equality across the eight bands at each skill value. The direct weight-band x baseline-skill interaction (p = 0.181) tested whether the complete pattern changed with skill.

**Figure 3.**
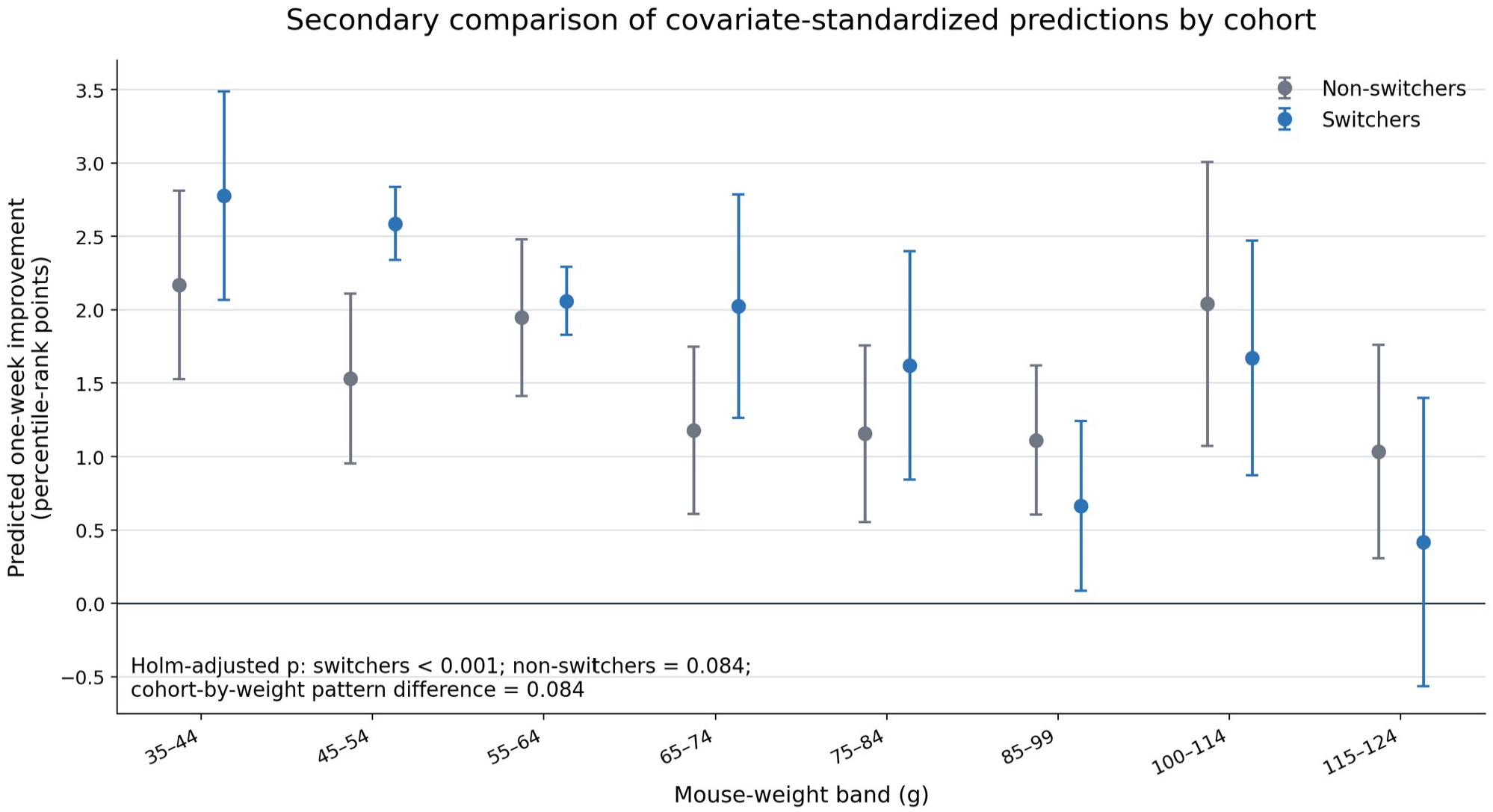
Secondary joint-model comparison of covariate-standardized predicted one-week percentile-rank improvement for switchers and non-switchers. Points show average model predictions for each cohort and weight band after standardizing over that cohort’s observed covariate distribution; error bars show pointwise 95% confidence intervals. Values are percentile-rank points. Across the three related joint-model tests, predicted improvement differed across bands for switchers (Holm-adjusted p < 0.001) but not for non-switchers (p = 0.084), and the complete eight-band patterns were not significantly different between cohorts (p = 0.084).

Eleven of the 28 overall band-pair comparisons had raw empirical permutation p < 0.05. Significant differences most often indicated greater predicted improvement for a band between 35 and 74 g than for 85-99 or 115-124 g; predicted improvement was also greater for 100-114 g than for 85-99 g. No weight band exceeded every other band, so the complete set of comparisons did not identify one uniquely optimal band. Raw empirical permutation results for the overall and skill-profile pairwise comparisons are reported in Table 3 and sections 3.3-3.5.

Raw empirical permutation results for the overall and skill-profile pairwise comparisons are reported in Table 3 and sections 3.3-3.5.

**Table 3.** Weight-band comparisons with raw empirical permutation p < 0.05 among switchers Each difference is e predicted one-week percentile-rank improvement for the first band minus that for the second band. Confidence intervals are covariance-aware, model-based 95% intervals; p values are two-sided raw empirical values from 1,000 weight-label permutations. Because the confidence intervals and permutation p values use different reference distributions, an interval can include zero even when the corresponding raw empirical p value is less than 0.05. All other pairwise comparisons had p ≥ 0.05.

| Skill profile | First weight band (g) | Second weight band (g) | Difference in predicted improvement (percentile-rank points) | 95% model-based CI | Raw permutation $p$ |
| --- | --- | --- | --- | --- | --- |
| Overall | 35-44 | 75-84 | 1.04 | -0.02 to 2.10 | 0.046 |
| Overall | 35-44 | 85-99 | 2.07 | 1.15 to 2.98 | 0.001 |
| Overall | 35-44 | 115-124 | 2.22 | 0.98 to 3.45 | 0.002 |
| Overall | 45-54 | 55-64 | 0.47 | 0.14 to 0.81 | 0.005 |
| Overall | 45-54 | 85-99 | 1.91 | 1.25 to 2.56 | 0.001 |
| Overall | 45-54 | 115-124 | 2.06 | 1.00 to 3.11 | 0.001 |
| Overall | 55-64 | 85-99 | 1.43 | 0.78 to 2.09 | 0.001 |
| Overall | 55-64 | 115-124 | 1.58 | 0.53 to 2.63 | 0.008 |
| Overall | 65-74 | 85-99 | 1.39 | 0.39 to 2.39 | 0.010 |
| Overall | 65-74 | 115-124 | 1.54 | 0.24 to 2.83 | 0.019 |
| Overall | 100-114 | 85-99 | 1.13 | 0.09 to 2.17 | 0.036 |
| Q25 | 35-44 | 85-99 | 2.48 | 1.05 to 3.90 | 0.004 |
| Q25 | 45-54 | 75-84 | 1.53 | 0.31 to 2.75 | 0.044 |
| Q25 | 45-54 | 85-99 | 2.41 | 1.51 to 3.31 | 0.001 |
| Q25 | 45-54 | 115-124 | 1.72 | 0.30 to 3.13 | 0.040 |
| Q25 | 55-64 | 85-99 | 1.96 | 1.10 to 2.82 | 0.001 |
| Median | 35-44 | 115-124 | 1.99 | 0.43 to 3.56 | 0.012 |
| Median | 45-54 | 85-99 | 0.92 | 0.03 to 1.82 | 0.031 |
| Median | 45-54 | 100-114 | 1.43 | 0.28 to 2.58 | 0.008 |
| Median | 45-54 | 115-124 | 2.27 | 0.90 to 3.65 | 0.001 |
| Median | 55-64 | 100-114 | 1.16 | 0.01 to 2.32 | 0.031 |
| Median | 55-64 | 115-124 | 2.01 | 0.63 to 3.38 | 0.005 |
| Median | 75-84 | 115-124 | 1.63 | -0.07 to 3.34 | 0.048 |
| Q75 | 35-44 | 115-124 | 1.97 | 0.10 to 3.83 | 0.019 |
| Q75 | 45-54 | 115-124 | 2.19 | 0.50 to 3.87 | 0.006 |
| Q75 | 55-64 | 115-124 | 1.95 | 0.26 to 3.65 | 0.018 |
| Q75 | 65-74 | 115-124 | 1.97 | -0.13 to 4.06 | 0.040 |
| Q75 | 75-84 | 115-124 | 2.08 | -0.05 to 4.22 | 0.029 |

**Table 4.**
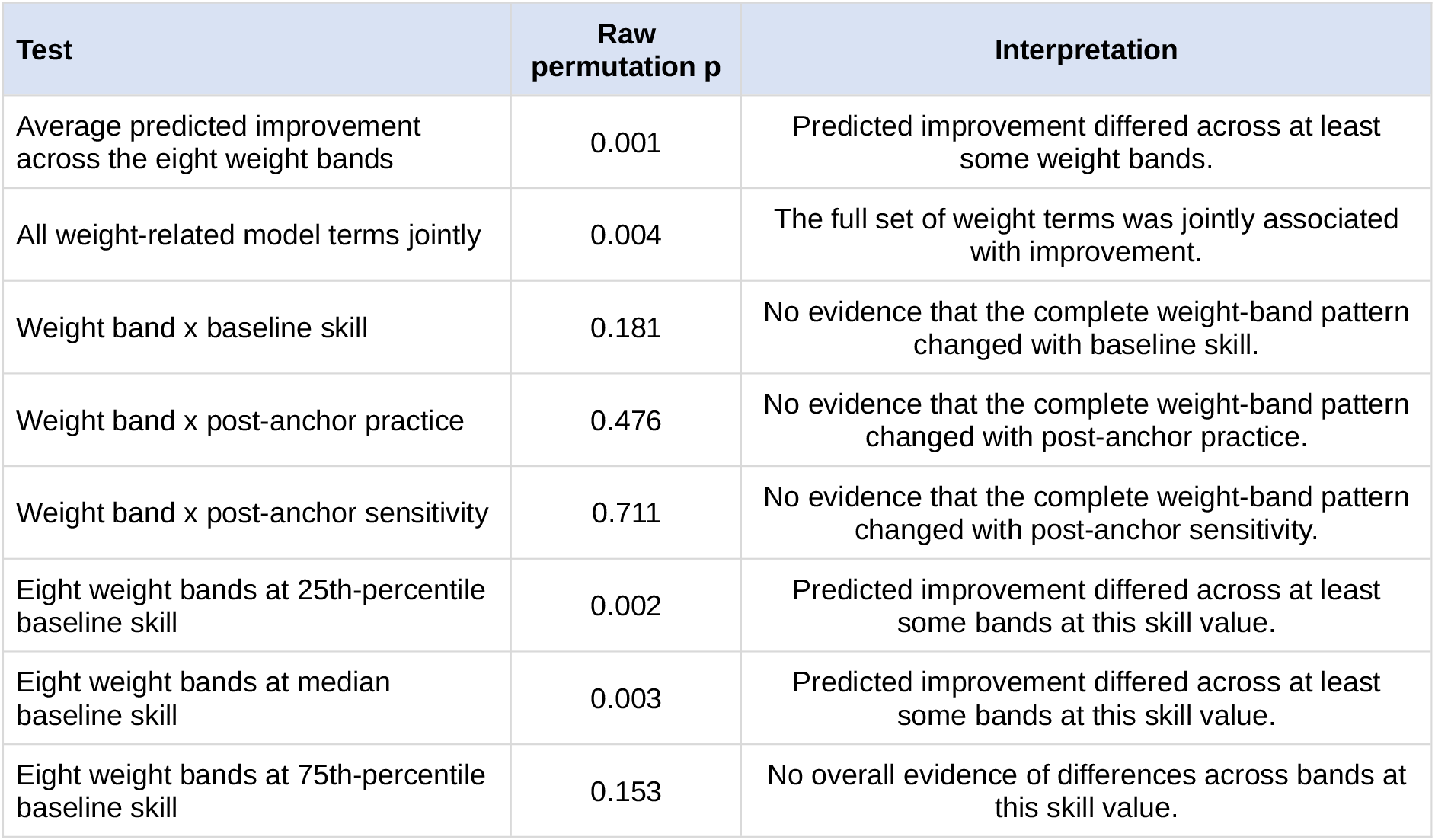
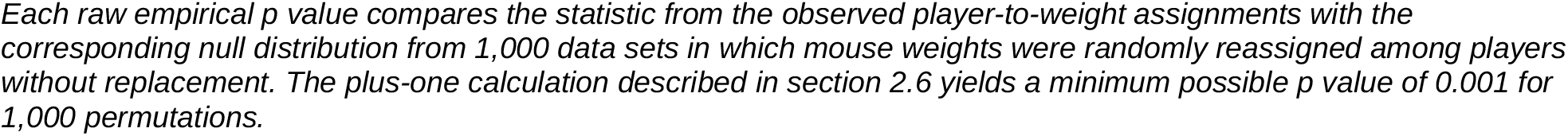
Weight-label permutation tests among switchers.

| Test | Raw permutation p | Interpretation |
| --- | --- | --- |
| Average predicted improvement across the eight weight bands | 0.001 | Predicted improvement differed across at least some weight bands. |
| All weight-related model terms jointly | 0.004 | The full set of weight terms was jointly associated with improvement. |
| Weight band × baseline skill | 0.181 | No evidence that the complete weight-band pattern changed with baseline skill. |
| Weight band × post-anchor practice | 0.476 | No evidence that the complete weight-band pattern changed with post-anchor practice. |
| Weight band × post-anchor sensitivity | 0.711 | No evidence that the complete weight-band pattern changed with post-anchor sensitivity. |
| Eight weight bands at 25th-percentile baseline skill | 0.002 | Predicted improvement differed across at least some bands at this skill value. |
| Eight weight bands at median baseline skill | 0.003 | Predicted improvement differed across at least some bands at this skill value. |
| Eight weight bands at 75th-percentile baseline skill | 0.153 | No overall evidence of differences across bands at this skill value. |

Each raw empirical $p$ value compares the statistic from the observed player-to-weight assignments with the corresponding null distribution from 1,000 data sets in which mouse weights were randomly reassigned among players without replacement. The plus-one calculation described in section 2.6 yields a minimum possible $p$ value of 0.001 for 1,000 permutations.

### 3.3 Overall pairwise comparisons

Using two-sided raw empirical permutation p values and a significance threshold of 0.05, 11 of the 28 overall pairwise contrasts were statistically significant. Predicted improvement was greater for 35-44 g than for 75-84, 85-99, and 115-124 g; for 45-54 g than for 55-64, 85-99, and 115-124 g; for 55-64 g than for 85-99 and 115-124 g; and for 65-74 g than for 85-99 and 115-124 g. Predicted improvement was also greater for 100-114 g than for 85-99 g. Differences ranged from 0.47 to 2.22 percentile-rank points (Table 3). The distribution of comparisons is consistent with a non-monotonic association rather than a sequence in which predicted improvement increases continuously as weight decreases.

### 3.4 Variation by baseline skill

The permutation omnibus test showed statistically significant differences in predicted improvement across the eight weight bands at the 25th-percentile baseline-skill value (raw empirical p = 0.002) and the median value (p = 0.003), but not at the 75th-percentile value (p = 0.153). At the 25th percentile, five pairwise comparisons had raw empirical p < 0.05: 35-44 versus 85-99 g; 45-54 versus 75-84, 85-99, and 115-124 g; and 55-64 versus 85-99 g. At the median, seven comparisons met this threshold: 35-44 versus 115-124 g; 45-54 versus 85-99, 100-114, and 115-124 g; 55-64 versus 100-114 and 115-124 g; and 75-84 versus 115-124 g. At the 75th percentile, five individual comparisons had raw empirical p < 0.05, each indicating greater predicted improvement for one of 35-44, 45-54, 55-64, 65-74, or 75-84 g than for 115-124 g (Table 3).

The profile-specific omnibus tests and the direct weight-band x baseline-skill interaction answer different questions. The profile-specific tests ask whether predicted improvement differs among the eight weight bands at one selected skill value. The interaction test asks whether those between-band differences themselves change as baseline skill changes. Predicted improvement differed across bands at the 25th-percentile and median skill values, but the interaction was not significant (raw empirical p = 0.181). In plain terms, some weight bands were associated with different predicted improvement at the selected lower and middle skill values, but the data did not show that players at different skill levels had statistically different weight improvement relationships or different optimal weights. The individual 75th-percentile comparisons are descriptive because the corresponding eight-band omnibus test was not significant.

### 3.5 Weight-label permutation results

All 1,000 permuted models converged. The omnibus test of differences in predicted improvement across the eight bands was statistically significant (raw empirical p = 0.001), as was the joint test of all weight-related model terms (p = 0.004). The interactions of weight band with baseline skill (p = 0.181), post-anchor practice (p = 0.476), and post-anchor mouse sensitivity (p = 0.711) were not statistically significant.

### *3*.*6* Secondary comparison with non-switchers

In the secondary joint model of 9,067 switchers and 4,991 non-switchers, the three related omnibus tests were treated as one family and evaluated with Holm correction. Predicted improvement differed across weight bands for switchers (Holm-adjusted p < 0.001). For non-switchers, the uncorrected test yielded p = 0.046, but the Holm-adjusted result was not statistically significant (p = 0.084). The complete eight-band pattern also did not differ significantly between switchers and non-switchers after correction (uncorrected p = 0.042; Holm-adjusted p = 0.084). Thus, the secondary comparison did not establish either an association between weight band and improvement during stable mouse use or a statistically different overall pattern between cohorts. The non-switcher results provide context for the primary switcher analysis but do not function as a randomized control comparison.

## 4. Discussion

### 4.1 Principal findings

The central finding is that post-switch mouse-weight band was statistically significantly associated with predicted one-week percentile-rank improvement among switchers. The largest predictions in player skill performance rank improvement occurred for switchers whose incoming mice weighed 35-44, followed by 45-54, and then 55-64 g, and the 11 significant overall pairwise comparisons most often indicated greater predicted improvement for bands from 35-74 g than for 85-99 or 115-124 g. The 1,000 weight-label-permutations supported differences in predicted improvement across the eight bands (raw empirical p = 0.001). The higher prediction for 100-114 g than for adjacent heavier bands and the absence of one uniformly superior band indicate that the association was non-monotonic; there is not a uniform “lighter is better” finding.

Predicted improvement differed across the eight bands at the 25th-percentile and median baseline-skill values (p = 0.002 and p = 0.003) but not at the 75th-percentile value (p = 0.153). These results show that some mouse weight bands differed at the selected lower and middle skill values. They do not show that the weight-improvement relationship itself changed with skill: the direct weight-band x baseline-skill interaction was not significant (p = 0.181). Therefore, the skill-profile results describe the fitted pattern at selected skill values but do not demonstrate different optimal weights for lower-, middle-, and higher-skill players.

The results do not identify one uniquely optimal mouse-weight band and should not be extrapolated below 35 g. No band was associated with significantly greater predicted improvement than every other band. The higher prediction at 100-114 g than at 85-99 or 115-124 g and the wider uncertainty in less-populated bands also argue against interpreting the results as a smooth relationship in which performance improvement increases continuously as mouse weight decreases.

### 4.2 Relation to prior work

The present study addresses a limitation identified in our earlier research, in which differences in player equipment were among the factors that were either unmeasured or not accounted for (Listman et al., 2021). Classical Human Computer Interface studies established that the design and control mapping of a pointing device can change movement time and error rate (Card et al., 1978; MacKenzie et al., 1991; Casiez et al., 2008), but those studies generally assigned a device for a brief controlled task. Here, the devices were not assigned by the investigators and were used during self-directed video game play over a period of one week. The naturalistic design provides scale and ecological validity, but not the experimental control needed to attribute the observed association to weight alone.

The controlled mouse-weight studies cited here enrolled 12-72 participants and generally measured immediate or same-session outcomes. Across small differences of 7-13 g, Li et al. (2019, 2022) and Wang et al. (2023) found preferences for lighter mice but little difference in several performance, muscle-activity, or kinematic measures. Across wider ranges, Yan et al. (2022) found the highest error rate at 40 g and the strongest preference near 60 g; Chen et al. (2012) found a non-monotonic relation between mouse weight and forearm muscular load; and Conroy et al. (2022) reported better target acquisition with 50-90 g than with 100 g. The present study extends this literature by evaluating one-week performance change after observed mouse-model changes in a much larger naturalistic cohort. Its comparatively high skill percentile rank improvement predictions for 35-44, 45-54, and 55-64 g and the frequent pairwise differences favoring bands from 35-74 g over 85-99 or 115-124 g are consistent with mouse mass being associated with performance, whereas the 100-114 g rebound and lack of one uniformly superior band are consistent with a non-monotonic relationship. Differences in design and outcome prevent treating these results as a direct replication of any single laboratory study.

Mouse sensitivity changes the mapping between hand displacement and view rotation (Casiez et al., 2008; Boudaoud et al., 2023), whereas baseline skill and practice distribution are associated with learning trajectories in large online samples (Stafford and Dewar, 2014; Listman et al., 2021). Sensitivity and practice were therefore treated as substantive covariates. Post-anchor practice was measured during the outcome week and could be both a determinant and a consequence of early post-switch performance; adjustment therefore addresses the selected conditional association rather than a total causal effect.

### 4.3 Motor adaptation as a possible mechanism

Two possible accounts of short-term performance change after a hardware switch are motor adaptation and an immediate performance advantage or disadvantage conferred by the new hardware. Motor adaptation recalibrates an existing skill to changed dynamics or sensorimotor mappings, whereas motor-skill acquisition typically develops over a longer period of practice (Haith and Krakauer, 2013; Diedrichsen and Kornysheva, 2015; Krakauer et al., 2019; Listman et al., 2021). A change in mouse mass alters the dynamics of the hand-held device even when software sensitivity remains constant. Related work has demonstrated perceptual-motor recalibration after a change in implement weight and anticipatory adjustment to changed object dynamics (Flanagan and Wing, 1997; Shadmehr and Mussa-Ivaldi, 1994; Scott and Gray, 2010).

Adaptation is not a single process with one time constant. Fast and slow components can differ in learning and retention (Smith et al., 2006), and re-exposure or repeated practice can affect explicit strategy and implicit recalibration differently (Avraham et al., 2021; Coltman et al., 2019; Huberdeau et al., 2019). Individuals can also differ in their use of explicit and implicit processes (Taylor et al., 2014; McDougle et al., 2016; McDougle and Taylor, 2019), and expertise can affect adjustment to a change in implement mass (Pan et al., 2024). These findings make adaptation a plausible framework for studying performance after a mouse switch. They do not show that adaptation caused the observed eight-band pattern, and the present data contain no direct measure of the adaptation process or of an immediate hardware advantage.

The non-switcher cohort provided a secondary estimate of weight-band associations over a comparable interval without an observed mouse change. Because a pseudo-anchor did not coincide with a physical-device change, non-switchers were not exposed to the same observable re-adaptation event as switchers. After correction across the three related joint-model tests, predicted improvement did not differ significantly across weight bands among non-switchers, and the complete eight-band patterns were not significantly different between cohorts. In practical terms, the comparison did not show a statistically reliable weight-band pattern during stable mouse use, but it also did not establish that the switcher and non-switcher patterns were different. The comparison supplies context for the primary switcher result; it cannot distinguish adaptation from an immediate hardware association and is not a randomized control analysis.

### 4.4 Practical interpretation

The complete pattern of predicted one-week improvement across all eight weight bands is more informative than a single target weight. In the primary switcher analysis, the largest predicted skill improvement occurred for 35-44, followed by 45-54, and then 55-64 g. The 11 statistically significant overall pairwise comparisons most often indicated greater predicted improvement for bands from 35-74 g than for 85-99 or 115-124 g, and 100-114 g also exceeded 85-99 g. However, no band exceeded every other band, and the non-monotonic pattern does not support a universal prescription for one mouse weight.

The selected baseline-skill profiles provide additional descriptions of the fitted eight-band pattern. Predicted improvement differed across bands at the 25th-percentile and median skill values, whereas the 75th-percentile omnibus test was not significant. Because the direct weight-band x baseline-skill interaction was also not significant, these results do not establish that the association between mouse weight and improvement differs by skill level. Individual comfort, fit, sensitivity, task demands, and adaptation time may also influence equipment choice and performance.

### 4.5 Strengths and limitations

Strengths include a large naturalistic switcher cohort, explicit handling of participant overlap, outcome-independent sensitivity-validity rules, complete cross-task calibration of analyzed performance observations, adjustment for practice intensity and mouse sensitivity, and reporting of all eight supported weight bands. The 1,000 weight-label permutations supplied empirical null distributions for the weight-related omnibus, interaction, profile-specific, and pairwise statistics. Covariate standardization placed each band’s predicted improvement on the same observed distribution of switcher baseline skill, practice, and sensitivity. Mouse-model identity was intentionally excluded from the inferential model so that weight remained the focal device characteristic; model identity nevertheless remained integral to exposure assignment and eligibility. The quota-sampled non-switcher cohort added a matched-time contextual comparison without defining the primary research question.

Several limitations constrain interpretation. Mouse-model changes and device weights were observed rather than assigned or randomized. Residual confounding may arise from player motivation to change gaming peripherals, concurrent changes in setup or play behavior, and physical mouse properties correlated with weight. The label-permutation analysis breaks the observed assignment of mouse weights to players, but it does not recreate randomized exposure, preserve every association between weight and measured player characteristics, or remove unmeasured confounding. It therefore provides a sensitivity analysis rather than a causal randomization test. To mitigate measured differences, the model included mouse sensitivity settings, pre-anchor date skill level, and practice intensity, which we have previously shown to be associated with skill improvement (Listman et al., 2021; Listman et al. unpublished results). The change-score outcome can also be affected by regression to the mean and measurement error, particularly when baseline values are extreme (Barnett et al., 2005; Vickers and Altman, 2001). Baseline-skill adjustment reduces but does not eliminate this concern.

Fixed bands sacrifice within-band resolution, although they avoid unsupported interpolation across gaps in commercially produced mouse weights. Sample sizes of upper-weight band cells were substantially smaller than the dominant 45-64 g cells, resulting in wider uncertainty for predicted values at upper-weight mouse bands. The one-week horizon does not establish whether associations in performance persist, reverse, or disappear over longer periods. Finally, the analysis concerns the included Aimlabs tasks and a calibrated-rank outcome, not all game genres or ergonomic outcomes.

The strict device-attribution requirement limits generalizability to mice whose manufacturer, model, and published mass could be resolved consistently. The final post-anchor exposure set nevertheless included 135 manufacturer-model combinations from 26 manufacturers.

The permutation analysis evaluated 28 overall and 84 skill-profile pairwise contrasts. To retain the full band-specific pattern, Table 3 reports every comparison with raw empirical p < 0.05; these results are interpreted together with the corresponding eight-band omnibus tests and the complete set of predicted values. With 1,000 permutations, empirical p values were resolved in increments of 1/1,001, with a minimum attainable value of 0.001.

## 5. Conclusions

The complete pattern of predicted one-week improvement across all eight weight bands is more informative than results for a single target weight. In this analysis of performance data from 9,067 Aimlabs switchers, the largest predictions of skill improvement occurred for switchers whose incoming mice weighed 35-44 g, followed by 45-54 g, then 55-64 g. The 11 statistically significant overall pairwise comparisons most often indicated greater predicted improvement for bands from 35-74 g than for 85-99 or 115-124 g, and 100-114 g predicted skill improvement also exceeded that of 85-99 g. However, no mouse weight band exceeded every other band, and the non-monotonic pattern does not support a universal prescription for one mouse weight.

The selected baseline-skill profiles provide additional descriptions of the fitted eight-band pattern. Predicted improvement differed across bands at the 25th-percentile and median skill values, whereas the 75th-percentile omnibus test was not significant. Because the direct weight-band x baseline-skill interaction was also not significant, these results do not establish that the association between mouse weight and improvement differs by skill level. Individual comfort, fit, sensitivity, task demands, and adaptation time may also influence equipment choice and performance.

## Declarations

### Author contributions

Jennifer B. Listman: Conceptualization, Methodology, Formal analysis, Visualization, Writing - original draft, and Writing - review and editing. Wayne E. Mackey: Conceptualization, Methodology, Software, Resources, and Writing - review and editing. Both authors read and approved the manuscript.

## Acknowledgments

The authors thank the Aimlabs players whose gameplay and performance data contributed to this work.

